# Nutrient-dependent mTORC1 signaling in coral-algal symbiosis

**DOI:** 10.1101/723312

**Authors:** Philipp A. Voss, Sebastian G. Gornik, Marie R. Jacobovitz, Sebastian Rupp, Melanie S. Dörr, Ira Maegele, Annika Guse

## Abstract

To coordinate development and growth with nutrient availability, animals must sense nutrients and acquire food from the environment once energy is depleted. A notable exception are reef-building corals that form a stable symbiosis with intracellular photosynthetic dinoflagellates (family *Symbiodiniaceae* (LaJeunesse et al., 2018)). Symbionts reside in ‘symbiosomes’ and transfer key nutrients to support nutrition and growth of their coral host in nutrient-poor environments (Muscatine, 1990; Yellowlees et al., 2008). To date, it is unclear how symbiont-provided nutrients are sensed to adapt host physiology to this endosymbiotic life-style. Here we use the symbiosis model *Exaiptasia pallida* (hereafter *Aiptasia*) to address this. *Aiptasia* larvae, similar to their coral relatives, are naturally non-symbiotic and phagocytose symbionts anew each generation into their endodermal cells (Bucher et al., 2016; Grawunder et al., 2015; Hambleton et al., 2014). Using cell-specific transcriptomics, we find that symbiosis establishment results in downregulation of various catabolic pathways, including autophagy in host cells. This metabolic switch is likely triggered by the highly-conserved mTORC1 (mechanistic target of rapamycin complex 1) signaling cascade, shown to integrate lysosomal nutrient abundance with animal development (Perera and Zoncu, 2016). Specifically, symbiosomes are LAMP1-positive and recruit mTORC1 kinase. In symbiotic anemones, mTORC1 signaling is elevated when compared to non-symbiotic animals, resembling a feeding response. Moreover, symbiosis establishment enhances lipid content and cell proliferation in *Aiptasia* larvae. Challenging the prevailing belief that symbiosomes are early arrested phagosomes (Mohamed et al., 2016), we propose a model in which symbiosomes functionally resemble lysosomes as core nutrient sensing and signaling hubs that have co-opted the evolutionary ancient mTORC1 pathway to promote growth in endosymbiotic cnidarians.

## Results and Discussion

### Physiological effects of symbiont uptake in *Aiptasia* larvae

Corals have extensive lipid stores to endure periods of low nutrient input (Stimson, 1987). Coral larvae depend on maternally deposited lipid droplets to fuel embryogenesis and development (Marlow and Martindale, 2007). We monitored the abundance of lipid droplets in *Aiptasia* larvae over time and found that lipid stores are plentiful 2 days post fertilization (dpf), largely consumed after 6 dpf and virtually depleted by 10 dpf (Figure 1A). Cell proliferation measured by EdU labeling decreases from 44% at 1 dpf to below 4% from 4 dpf onwards (Figure 1B and C). This suggests that after a few days, aposymbiotic larvae have consumed pre-deposited energy reserves and enter a stationary phase until alternative nutrient sources or other environmental cues initiate further development and metamorphosis (Bucher et al., 2016). Biogenesis of lipid droplets in coral cells depends on the symbionts (Chen et al., 2012; Muscatine et al., 1994). To test the effects of symbiosis establishment on changes in lipid content and cell proliferation in *Aiptasia* larvae, we quantified the number of lipid droplets at various times after symbiont uptake. At 10 dpf, symbiotic larvae contain more lipid droplets 2 and 8 days after symbiont uptake when compared to aposymbiotic larvae (Figure 1D and E). Moreover, we found that 3 days after infection, cell proliferation was significantly higher in symbiotic larvae (adjusted p-value < 0.001) when compared with uninfected control larvae (Figure 1F). This suggests that symbiosis establishment has positive effects on *Aiptasia* larval physiology, most likely due to an improved nutritional status through the initiated endosymbiotic nutrient transfer, similar to what has been observed in coral larvae (Kopp et al., 2016). Thus, symbiosis establishment in *Aiptasia* larvae provides a suitable framework to analyze the molecular mechanisms of how symbiont-derived nutrients are sensed and integrated into host metabolism.

**Figure 1.**
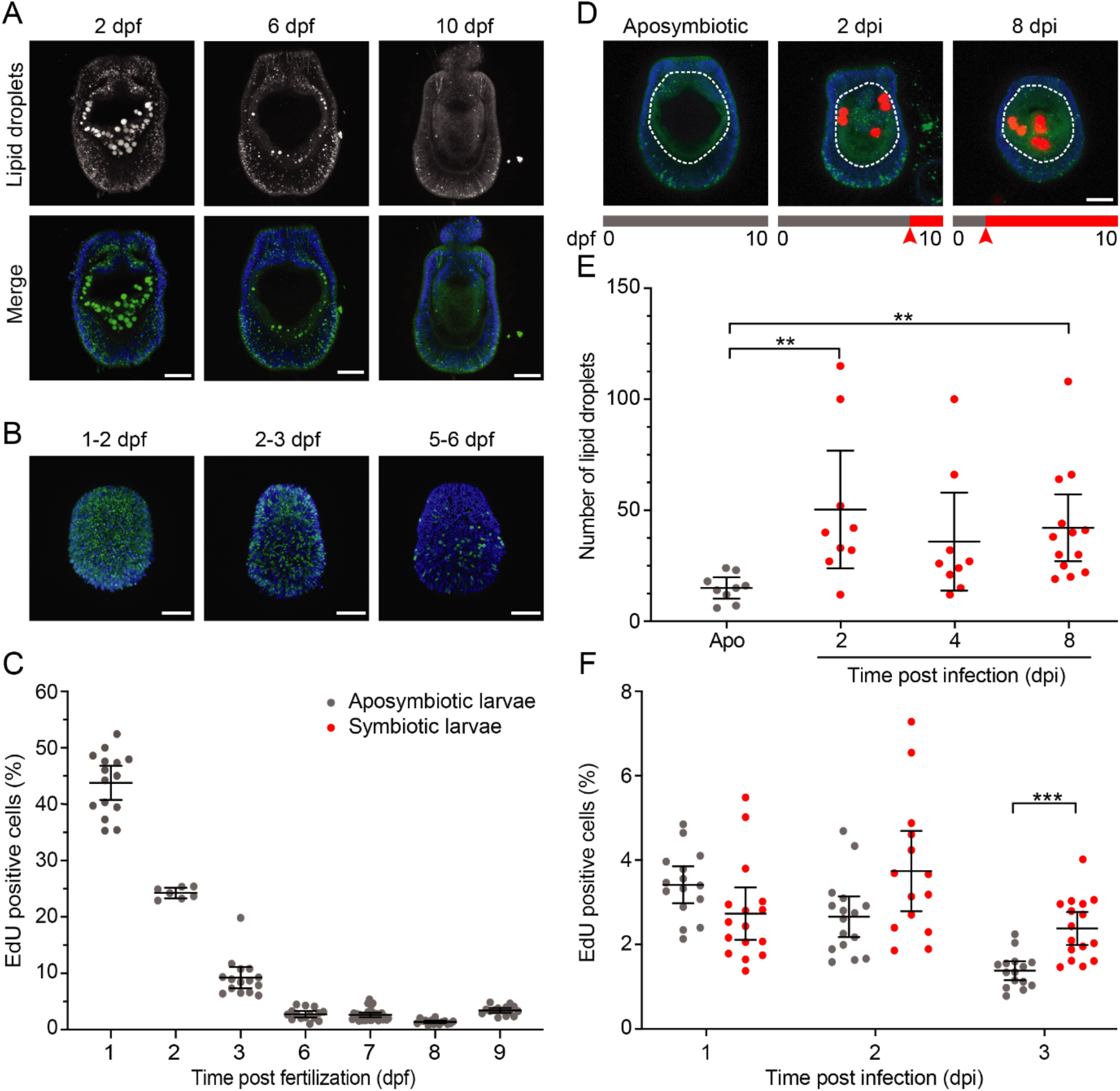
Physiological effects of symbiosis establishment on *Aiptasia* larvae. **A)** Representative images of *Aiptasia* larvae 2, 6, and 10 days post fertilization (dpf) stained with the lipid dye Nile Red. Size and number of lipid droplets decrease with the age of larvae. Images are maximum projections of 30 μm thick larval mid-sections. Colors in merge are nuclei in blue (Hoechst 33258) and lipid droplets in green (Nile Red); scale bars represent 25 μm. **B)** Representative images of *Aiptasia* larvae 2, 3, and 6 dpf, each after 24 h EdU pulse used in the cell proliferation assay in Figure 1C. Images are maximum projections of whole larvae. Colors in merge are nuclei in blue (Hoechst 33258) and nuclei of proliferating cells in green (EdU); scale bars represent 25 μm. **C)** Cell proliferation in aposymbiotic larvae measured by the proportion of nuclei that had incorporated EdU in the preceding 18 h (1dpf) or 24 h (2 – 9 dpf). Each data point represents one larva (n = 7 - 16). Error bars represent mean and 95 % confidence interval **D)** Representative images of *Aiptasia* larvae 10 dpf, that were either aposymbiotic or infected 2 or 8 days prior to staining with Nile Red used in Figure 1E. Images are maximum projections of 30 μm thick larval mid-sections. Colors in merge are nuclei in blue (Hoechst 33258), lipid droplets in green (Nile Red), and symbiont autofluorescence in red; scale bars represent 25 μm. **E)** Comparison of number of lipid droplets in aposymbiotic (gray) and symbiotic larvae (red) at 2, 4, and 8 days after symbiont uptake. Each data point represents one larva (n = 9 - 13). Error bars represent mean and 95 % confidence interval. For ImageJ macros used for quantification, see File S1 and File S2 **F)** Comparison of cell proliferation in aposymbiotic (gray) and symbiotic (red) larvae 1, 2 or 3 days post infection. Each data point represents 1 larva (n = 15 - 16). Error bars represent mean and 95 % confidence interval

### Host cell gene expression is down-regulated in response to symbiont uptake

To monitor the transcriptional changes in host cells induced by symbiont acquisition, we infected *Aiptasia* larvae 6-7 dpf for 24-48 h, dissociated both symbiotic and aposymbiotic control larvae into symbiotic (red circle) and aposymbiotic (gray circle) cells (Figure 2A and B). We then collected pools of 7-20 individual symbiotic and aposymbiotic cells (Figure 2B, center panels) for comparison by RNA-Seq. This set of samples allows analysis of gene expression in symbiotic and aposymbiotic cells shortly after symbiont uptake with cell-type-specific resolution. This approach is clearly distinct from transcriptomic studies comparing gene expression of whole individuals of anemone or coral species (Lehnert et al., 2014; Matthews et al., 2017; Mohamed et al., 2016; Wolfowicz et al., 2016; Yuyama et al., 2018) and is a vital prerequisite for understanding the effects of symbiont uptake at the cellular level.

**Figure 2.**
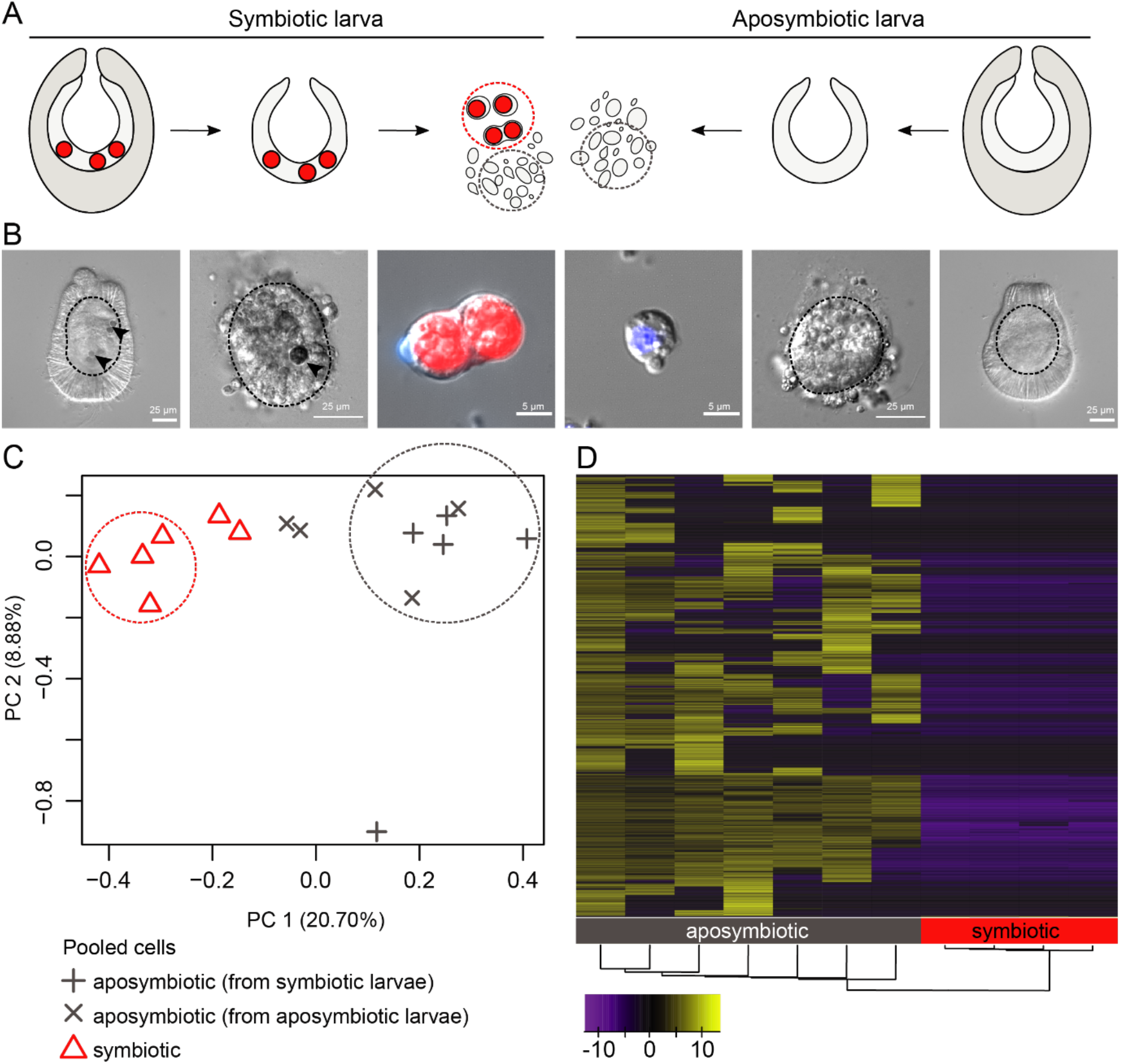
Host cell gene expression is down-regulated in response to symbiont uptake. **A)** Sampling approach for generation of cell-type-specific transcriptomes. Symbiotic larvae were dissociated using Pronase, yielding naked endodermal tissue. Maceration resulted in dissociated cells, of which pools of 7 to 20 cells of either symbiotic (red dashed circles) or aposymbiotic cells (gray dashed circles) were isolated for transcriptomic analysis. **B)** Representative DIC images of dissociation procedure as in Figure 2A. Arrowheads indicate symbionts. Dotted circles represent endodermal tissue. Colors in merge are nuclei in blue (Hoechst 33258), symbiont autofluorescence in red, and DIC in gray. **C)** Principal Component Analysis (PCA) plot of host gene expression for all genes in all replicates. Dotted circles indicate the samples used for further analysis of differential gene expression of aposymbiotic (gray) and symbiotic cells (red). **D)** Heatmap showing differentially expressed genes between symbiotic (red) and aposymbiotic cells (gray). 4,456 out of 27,334 genes were differentially expressed; 4,421 genes were down-regulated in the symbiotic condition and 35 genes were up-regulated (log2-fold change > 2; false-discovery-rate < 0.01). For a complete list of differentially expressed genes, see Table S1. For results of KEGG enrichment analysis, see Tables 1 and S2. For a schematic summarizing gene expression of core metabolic pathways, see Figure S1.

Principal component analysis (PCA) revealed considerable differences in gene expression between symbiotic and aposymbiotic endodermal cells. However, whether aposymbiotic cells originated from symbiotic or aposymbiotic larvae was indistinguishable (Figure 2C). This suggests that within the first 24-48 hours post infection (hpi), symbiont uptake influences gene expression in their respective host cells, yet does not elicit major effects in aposymbiotic cells within the same larva; thus highlighting the importance of our cell-type-specific approach. We next focused on a direct comparison of gene expression of the replicates most representative of the symbiotic (Figure 2C, red circle) and aposymbiotic state (Figure 2C, gray circle). We found 4,456 out of 27,334 genes to be differentially expressed (DEGs) (log2-fold change > 2; false-discovery-rate < 0.01). More than 99% of DEGs are down-regulated in the symbiotic state (4,421 genes vs. 35 up-regulated genes, Figure 2D, Table S1). A higher proportion of down-regulated genes in symbiotic animals has been previously reported in both anemones and corals (Matthews et al., 2017; Mohamed et al., 2016; Wolfowicz et al., 2016); however, the effect is much more pronounced in our dataset, likely due to the higher signal-to-noise ratio of transcripts from symbiotic vs. aposymbiotic cells.

### Host cell metabolism and autophagy are down-regulated upon symbiosis establishment

Using KEGG Enrichment analysis, we found that three cellular pathways (endocytosis, autophagy (yeast), peroxisome) were down-regulated in symbiotic cells (Table 1, Table S2). Expression of approximately 40 % (37/96) of the endocytosis genes and > 20% of autophagy genes including the key players ATG2, ATG4, ATG9, ATG13, ATG18, HIF1α, TSC1, PKCδ and RagC/D is suppressed, indicating that autophagy and endocytosis levels significantly drop upon symbiosis establishment. Moreover, various pathways of the KEGG categories metabolism, genetic information processing, and environmental information processing are enriched among the genes that were down-regulated in symbiotic cells (Table 1, Table S2). A detailed analysis of various core metabolic pathways revealed reduced gene expression of key enzymes, sometimes even below the detection limit (Figure S1). This suggests that numerous central pathways involved in sugar, lipid, and amino acid metabolism are down-regulated upon symbiont uptake (Table 1, Table S2, Figure S1). The affected KEGG pathways are either catabolic (incl. autophagy, peroxisomes, as well as starch and sucrose metabolism) or could be secondarily affected by the lack of primary metabolites, which would otherwise be provided by the now down-regulated catabolic pathways. This indicates that aposymbiotic host cells were metabolically active, but symbiont uptake induced a metabolic switch. Specifically, we speculate that in the absence of exogenous food, aposymbiotic *Aiptasia* larvae 6-7 dpf maintain homeostasis by autophagy. It appears that host cell metabolism is reprogrammed in response to symbiont uptake, potentially due to the initiation of nutrient transfer from symbionts to host cells.

**Table 1.** Enriched KEGG pathways among the down-regulated genes in symbiotic cells (p-value ≤0.15). For a complete list of all differentially expressed genes of each pathway, see supplementary Table S2.

| Pathway | KEGG Pathway ID |
| --- | --- |
| Metabolism |  |
| Synthesis and degradation of ketone bodies | [PATH:ko00072] |
| Terpenoid backbone biosynthesis | [PATH:ko00900] |
| Arginine biosynthesis | [PATH:ko00220] |
| Starch and sucrose metabolism | [PATH:ko00500] |
| Galactose metabolism | [PATH:ko00052] |
| N-Glycan biosynthesis | [PATH:ko00510] |
| Fructose and mannose metabolism | [PATH:ko00051] |
| Pyrimidine metabolism | [PATH:ko00240] |
| Glycosylphosphatidylinositol (GPI)-anchor biosynthesis | [PATH:ko00563] |
| Cellular Processes |  |
| Autophagy - yeast | [PATH:ko04138] |
| Peroxisome | [PATH:ko04146] |
| Endocytosis | [PATH:ko04144] |
| Genetic Information Processing |  |
| Basal transcription factors | [PATH:ko03022] |
| RNA polymerase | [PATH:ko03020] |
| RNA degradation | [PATH:ko03018] |
| Protein export | [PATH:ko03060] |
| RNA transport | [PATH:ko03013] |
| Non-homologous end-joining | [PATH:ko03450] |
| Spliceosome | [PATH:ko03040] |
| Ribosome biogenesis in eukaryotes | [PATH:ko03008] |
| Environmental Information Processing |  |
| Phosphatidylinositol signaling system | [PATH:ko04070] |

### Regulation of autophagy by MITF-like transcription factors is evolutionarily conserved

The switch between autophagy and biosynthesis in response to nutrient levels has been analyzed in detail in mammalian cells (Perera and Zoncu, 2016). When nutrients are scarce, the transcription factor EB (TFEB) induces autophagy and lysosomal biogenesis in a coordinated fashion by activating the transcription of ‘CLEAR (Coordinated Lysosomal Expression And Regulation) network’ genes (Settembre et al., 2011). TFEB binds to so-called ‘CLEAR elements’ (GTCACGTGAC), which are expanded E-box motifs (CANNTG), and additionally, induces an auto-regulatory feedback loop (Sardiello et al., 2009; Settembre et al., 2013). TFEB belongs to the MITF-family of transcription factors, which expanded into multiple transcription factors, including TFEB in vertebrates (Bouché et al., 2016; Sardiello, 2016). We identified a MITF-like protein (XP_020895872.1) in *Aiptasia* and conducted a phylogenetic analysis of MITF-family transcription factors across vertebrates and invertebrates (Figure 3A). The phylogeny recapitulates the previously reported MITF distribution including a vertebrate expansion of the MITF-family from one ancestral, invertebrate MITF-like gene. Amongst the invertebrate MITF-like TFs the *C. elegans* homolog has been shown to have a similar function as vertebrate TFEB, recognizing E-box motifs (CANNTG) in the promoters of its target genes to induce autophagy (Lapierre et al., 2013; O’Rourke and Ruvkun, 2013; Settembre et al., 2013). Similarly, we identified E-box motifs in the promoters of 16 out of 17 *Aiptasia* homologs of previously characterized mammalian TFEB target genes, including the MITF-like transcription factor itself (Figure 3B). For the *Aiptasia* MITF-like and 11 others of these genes, gene expression is significantly down-regulated in symbiotic cells (Figure 3B). This suggests that the coordinated regulation of autophagy and lysosomal biogenesis is the ancestral role of the MITF-like TF family in invertebrates, which may be used to adapt the cellular metabolism of host cells to incoming symbiont-derived nutrients.

**Figure 3.**
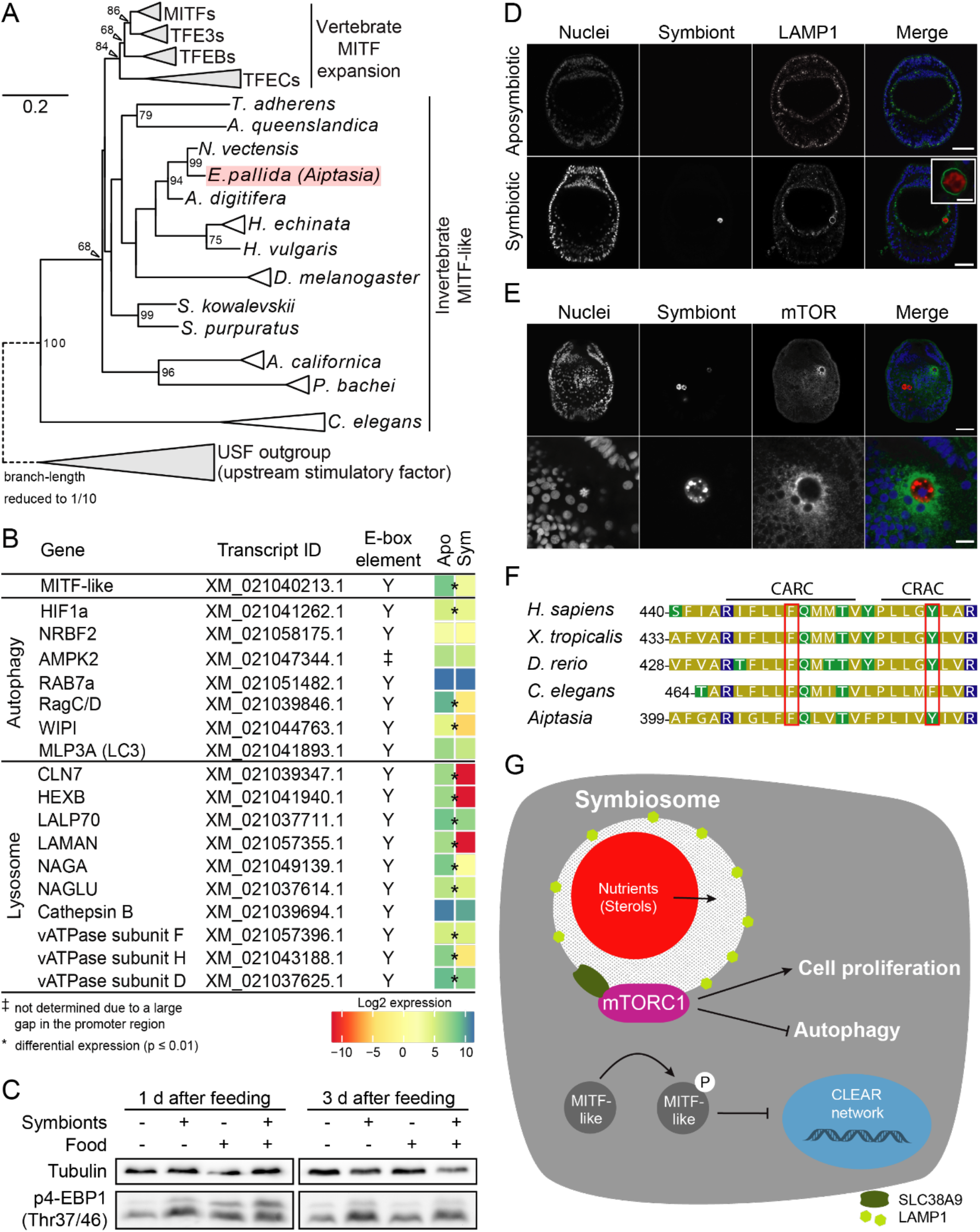
Symbiosis activates mTORC1 signaling. **A)** Maximum Likelihood phylogeny (PhyML) of MITF-family proteins across vertebrates and invertebrates. Only bootstrap values (ML percentage) ≥ 50 are shown. We used upstream regulatory factors, which are sister to the MITF-like TFs within the bHLH (basic helix-loop-helix) TFs, as an outgroup. Species names are italicized. Triangles represent collapsed branches. The position of *Aiptasia* is highlighted in light red. For raw sequences, trimmed alignments, and tree information, see File S3. For alignment of *Aiptasia* MITF-like and *Homo sapiens* TFEB, see Figure S2A. **B)** Overview of gene expression of *Aiptasia* homologs of CLEAR network genes involved in autophagy and lysosomes. Presence of E-box motifs in the promoter region of a gene is marked with “Y”. Asterisks “*” indicate differentially expressed genes (log2-fold change > 2; false-discovery-rate < 0.01). **C)** Representative Western blots of phosphorylated 4-EBP1 (Thr37/46) (p4-EBP1) comparing aposymbiotic and symbiotic polyps with (feeding 3 times per week, for > 3 weeks) and without food (no feeding for ≥ 3 weeks). For representative images of lipids in polyp macerates see Figure S2B. **D)** Representative images of LAMP1-immunofluorescence analysis of *Aiptasia*-LAMP1 (see also Figure S2C) in aposymbiotic and symbiotic larvae 6 dpf. Colors in merge are nuclei in blue (Hoechst 33258), LAMP1 in green and LAMP1 detected with Alexa488-anti-rabbit IgG, and symbiont autofluorescence in red; scale bars represent 25 μm for overviews and 5 μm for inset. **E)** Representative images of mTOR-immunofluorescence analysis occurring in ~50% of symbiotic larvae, 6 dpf. Colors in merge are nuclei in blue (Hoechst 33258), mTOR in green and symbiont autofluorescence in red; scale bars represent 25 μm for whole larva images (upper panels) and 5 μm for close up (lower panels). **F)** Protein sequence alignment of transmembrane domain 8 of SLC38A9 homologs. Key conserved phenylalanine (F) and tyrosine (Y) residues within the CARC and CRAC motifs boxed in red. **G)** Model of mTORC1 signaling to coordinate nutrient input by symbionts with host physiology in cnidarian endosymbiosis.

### Symbiosis activates mTORC1 signaling

In mammals, TFEB activity is controlled by phosphorylation of two serine residues by mTOR kinase, which are conserved in the *Aiptasia* MITF-like protein (Figure S2A) (Martina et al., 2012; Roczniak-Ferguson et al., 2012; Settembre et al., 2012). In the presence of nutrients, mTOR kinase is recruited as a member of mTORC1 to the lysosomal surface and is activated (Sancak et al., 2008). Phosphorylation leads to retention of TFEB in the cytoplasm leading to reduced expression of CLEAR network genes and thus down-regulation of autophagy (Martina et al., 2012; Roczniak-Ferguson et al., 2012; Settembre et al., 2012). We identified homologs of all major components of the mTORC1 complex, including mTOR kinase, raptor, deptor, and mLST8, except for PRAS40 (data not shown) in the *Aiptasia* genome. Two main mTOR kinase targets, S6 kinase 1 (S6K1) and eukaryotic translation initiation factor 4E (eIF4E)-binding protein 1 (4E-BP1), both of which stimulate protein synthesis (Hay and Sonenberg, 2004), are likewise conserved in *Aiptasia* (data not shown). To directly test whether the presence of endosymbionts activates mTOR kinase in *Aiptasia* adult anemones, we compared the levels of phosphorylated 4E-BP1 (p4-EBP1), a commonly used marker for mTOR activity, between aposymbiotic and symbiotic animals, with and without feeding. We find that p4E-BP1 levels are relatively low in starved, aposymbiotic animals, but increase upon feeding (Figure 3C, left panel). In comparison to well-fed aposymbiotic anemones, p4E-BP1 levels are equally high in symbiotic anemones without further increase upon additional feeding (Figure 3C, left panel).Interestingly, the increase of p4E-BP1 levels in aposymbiotic anemones after feeding disappeared within 3 days (Figure 3C, right panel). This suggests that the presence of symbionts activates mTOR kinase similarly to nutrient input by feeding. However, the effect of symbiosis is continuous, while nutrient input has to occur on a regular basis to maintain elevated mTORC1 signaling. In line with the role of mTORC1 signaling in promoting anabolic processes (Perera and Zoncu, 2016), tissue macerates of symbiotic anemones that have not been fed for 9 days contained prominent lipid droplets while aposymbiotic animals did not (Figure S2B). Regular feeding (3 times per week for 10 days) of aposymbiotic animals also promotes lipid droplet formation, albeit to a lesser extent than continuous nutrient input by endosymbiosis (Figure S2B). This is consistent with previous work showing that presence of symbionts is beneficial for *Aiptasia* promoting growth and asexual reproduction (Clayton and Lasker, 1985).

### Symbiosomes resemble lysosomes as nutrient-sensing and signaling centers

How do symbiont-derived nutrients activate mTORC1 in symbiotic cells? In mammalian cells, lysosomes play a key role to dynamically adapt cellular metabolism to nutrient availability (Perera and Zoncu, 2016). Abundant nutrient levels within the lysosomal lumen are sensed by the transmembrane protein SCL38A9, leading to mTORC1 recruitment to the lysosomal surface and signaling induction to promote biosynthesis and proliferation (Castellano et al., 2017). In the absence of lysosomal nutrients, lack of mTORC1 signaling drives the cell towards catabolism, including autophagy, to maintain homeostasis. To ask whether symbiosomes resemble lysosomes in their function as nutrient-sensing hubs, we first generated an *Aiptasia specific* LAMP1 (Lysosome-associated membrane protein 1) antibody (Figure S2C). LAMP1 is a highly glycosylated protein present on late endosomes and lysosomes, which provides structural integrity by forming a continuous carbohydrate lining on the inner leaflet of the lysosomal membrane (Eskelinen, 2006). Using immunofluorescence, we find that all symbiosomes are prominently decorated with LAMP1 (Fig. 3D). In accordance with being bona-fide signaling centers, individual symbiosomes are associated with mTOR kinase (Fig. 3E). This suggests that symbiosomes are lysosomal-like organelles with the capacity to function as mTORC1 signaling platforms in response to symbiosis. However, we noted that only ~50 % of symbiotic larvae at 6 dpf have mTOR-positive symbiosomes. This suggests that signaling activity may be dynamically regulated, for example, based on current photosynthesis and nutrient transfer rates.

In mammals, mTORC1 signaling is induced by elevated lysosomal cholesterol levels sensed by SCL38A9 (Castellano et al., 2017). Interestingly, corals and anemones are sterol auxotroph and receive the bulk of their sterols from their symbionts (Baumgarten et al., 2015; Goad L. J., 1981; Hambleton et al., 2019). Both have expanded their repertoire of NPC2 proteins, highly conserved lysosomal cholesterol binders that localize to the symbiosome and directly bind symbiont-derived sterols (Hambleton et al., 2019). Thus, such sterols may well be among the key nutrients that are sensed and trigger a metabolic shift in host cells by mTORC1 signaling. Accordingly, we found an SLC38A9 homolog in *Aiptasia* (transcript ID XM_021036759.2) containing the highly conserved cholesterol-responsive CARC and CRAC motifs previously identified in its mammalian homolog (Fig. 3F). Thus, the ability to measure sterol levels in lysosomes/symbiosomes to switch mTORC1-mediated nutrient-signaling on and off may have been co-opted in endosymbiotic cnidarians to adapt host metabolism to nutritional input by their photosynthetic symbionts.

## Conclusion

Taken together, we propose a model where symbiont-derived nutrients, such as sterols, are sensed in the lumen of the symbiosome, triggering mTORC1 recruitment and signaling. mTORC1 signaling leads to increased cell proliferation and, via the conserved MITF-like TF, to reduced autophagy levels as a response to symbiont uptake. More broadly, symbiosomes functionally resemble mammalian lysosomes as signaling hubs that coordinate nutrient abundance with proliferation and growth (Fig. 3G). This model challenges the hypothesis that the symbiosome is an arrested early phagosome (Davy et al., 2012; Mohamed et al., 2016). Based on observations that symbionts appear to be associated with the early endosomal marker Rab5, but devoid of the late endosomal/lysosomal marker Rab7 (Chen et al., 2004), it was thought that symbiosomes avoid fusion with the lysosome to allow intracellular persistence in host cells, a strategy that is employed by some intracellular pathogens such as *Streptococcus pyogenes* (Smith and May, 2013). However, other experimental evidence suggests that symbiosomes are acidified by a lysosomal vATPase and contain NPC2 proteins, both lysosomal proteins, contradicting the idea that interference with phagosomal maturation is key to symbiosome formation (Barott et al., 2015; Hambleton et al., 2019). In the future, it will be crucial to further characterize the symbiosome protein composition and its physiology to better understand how it is regulated to adapt symbiosome function to both host and symbiont. The ability to switch between biosynthetic and catabolic pathways to maintain cellular functions, even when nutrients are limited, is key for animal adaptation and evolution (Chantranupong et al., 2015). Our findings have important implications for understanding the mechanism and evolution of nutrient sensing in coral endosymbiosis, the cornerstone of coral-reef ecosystems, which are increasingly threatened by the disruption of the symbiosis (‘coral bleaching’) due to global warming.

## Acknowledgements

We thank Dinko Pavlinic and Vladimir Benes (Genecore Facility, EMBL Heidelberg) for assistance with the SmartSeq2 protocol and sequencing library preparation; David Ibberson (Deepseqlab, Heidelberg University) for assistance with the SmartSeq2 protocol; Carsten Rippe for access to the BioAnalyzer; Carlo Beretta (Math-Clinic, Heidelberg University) for help with image analysis; Jan Siemens, Shiying Lu and Jörg Pohle for advice on cell picking; Thomas Holstein and Steffen Lemke for advice and comments; Elizabeth A. Hambleton and Carmine Settembre for comments on the manuscript.

Funding was provided by the Deutsche Forschungsgemeinschaft (DFG) (Emmy Noether Program Grant GU 1128/3–1), the European Commission Seventh Framework Marie-Curie Actions (FP7-PEOPLE-2013-CIG), the H2020 European Research Council (ERC Consolidator Grant 724715) and a PhD scholarship within the Graduate School “Evolutionary Novelty & Adaptation by the Baden-Württemberg Landesgraduiertenförderung Program to PAV; and to SR by the CellNetworks Excellence Cluster (Heidelberg University) Postdoctoral Program.

## Author Contributions

Conceptualization, P.A.V. and A.G.; Methodology, P.A.V., S.G.G., A.G.; Software, P.A.V. and S.G.G.; Formal Analysis, P.A.V., S.G.G., S.R.; Investigation, P.A.V., S.G.G., M.R.J., S.R., M.S.D. and I.M.; Resources, A.G.; Data Curation, P.A.V. and S.G.G.; Writing - Original Draft, P.A.V. and A.G., Writing – Review & Editing, P.A.V., S.G.G., M.R.J., S.R., I.M. and A.G.; Visualization; P.A.V., S.G.G., M.R.J. and S.R.; Supervision, A.G., Project Administration, P.A.V. and A.G.; Funding Acquisition, A.G.

## Methods

### Live Organism Culture and Maintenance

#### Algal maintenance

For infection experiments of *Aiptasia* larvae, we used the clonal axenic culture of *Breviolum minutum* (GenBank Accession MK692539, hereafter referred to as SSB01). Cultures were grown in cell culture flasks in 0.22 μm filter-sterilized 1X Daigo’s IMK medium (398-01333, Nihon Pharmaceutical Co. Ltd.) on a 12h light:12h dark (12L:12D) cycle under 20–25 μmol m−2 s−1 of photosynthetically active radiation (PAR) at 26 °C.

#### Aiptasia culture conditions, spawning induction and larval culture conditions

*Aiptasia* clonal lines F003 and CC7 were maintained at 26 °C in a 12L:12D cycle. Animals were induced to spawn following the previously described protocol (Grawunder et al., 2015). *Aiptasia* larvae were maintained at ~300 larvae per ml in glass beakers in 0.22 μm filter-sterilized artificial sea water (FASW) at 26°C and exposed to a 12L:12D cycle.

### Transcriptome sample preparation

*Aiptasia* larvae (~300 per ml) were infected 6 or 7 dpf with 10^5^ SSB01 cells per ml for 24 or 48 hours or left aposymbiotic. A total of 3 to 5 infected larvae were transferred in 2 μl FASW to 5 ml of Calcium- and Magnesium-free artificial sea water (CMF-SW, doi:10.1101/pdb.rec12053). After incubation for 5 min, larvae were transferred to a 70 μl drop of Pronase (0.5 % in CMF-SW; 10165921001, Sigma-Aldrich Co. LLC) and sodium thio glycolate (STG, 1 % in CMF-SW; T0632, Sigma-Aldrich Co. LLC) on a glass microscopy slide. After mixing larvae by pipetting up and down in 20 μl 3 to 5 times, larvae were incubated for ca. 2 min until ectodermal cells began to be released from the larval body. The endodermal core was transferred to a drop of 70 μl FASW and remaining ectodermal cells were washed off by pipetting up and down five to 10 times in 20 μl. Endodermal tissue was transferred to a 70 μl drop of FASW on a cover slip and crushed using the tip of tweezers to yield single cells. The total time from beginning of dissociation of larvae to lysis was 30 min.

Pools of 7 to 20, either aposymbiotic or symbiotic, cells were picked using special microcapillary needles with openings of 8 to 12 μm diameter pulled with a P-97 Flaming/Brown Micropipette puller from glass capillaries (Science Products GB100T-8P). Glass capillaries were pre-loaded with 4.3 μl of lysis buffer (0.2 % TritonX-100, 1U/μl Protector RNase inhibitor (3335399001, Sigma-Aldrich Co. LLC), 1.25 μM oligo-dT30VN, and 2.5 mM dNTP mix) and collected cells were flushed out of the capillary with the lysis buffer into a PCR tube before flash-freezing in liquid nitrogen. After cell capture and lysis as described above, sequencing libraries were prepared as previously described (Picelli et al., 2014). Briefly, RNA was reverse transcribed, followed by pre-amplification of cDNA over 21 PCR cycles. cDNA libraries were then prepared for Illumina sequencing and sequenced on NextSeq500 with 75 bp paired-end sequencing.

### Computational Methods

#### Differential gene expression analysis

To exclude contaminations and, where applicable, symbiont-derived reads, paired-end reads were mapped to the *Aiptasia* genome version GCF_001417965.1 using HISAT2 version 2.1.0 at default settings, except - X 2000 --no-discordant --no-unal --no-mixed. Transcripts were quantified in Trinity v2.5.1 using salmon v0.10.2 at default settings using filtered reads. Principal component analysis was conducted using perl scripts supplied with Trinity for all samples. Differentially expressed transcripts between clustered symbiotic and aposymbiotic samples (circled in Fig. 2C) were detected using DEseq2 (log2-fold change > 2; false-discovery-rate < 0.01).

#### KEGG pathway enrichment analysis

Differentially expressed genes were examined for enrichment of KEGG pathway terms in R v3.4.1 using the ‘enricher’ function from the R package ‘clusterProfiler’ (Yu et al., 2012) at standard settings, visualized using the R package ‘ComplexHeatmap’ (Gu et al., 2016) and finalized using Adobe Illustrator CC 2018.

#### Phylogeny of MITF-family

For MITF phylogenies, metazoan MITF and USF (upstream stimulatory factor) homologs were identified from public databases. Alignments were generated using ClustalW v2.0 (Larkin et al., 2007) and manually corrected in Geneious v10.2.6 (Biomatters). Ambiguous sites and poorly aligned regions were removed automatically using trimAI set to ‘-automated1’ (Capella-Gutiérrez et al., 2009). We then determined the best-fitting substitution model using ModelFinder (set to ‘-m MF -msub nuclear’) within iqTree 1.6.10 and PROTTEST3 (set to ‘-JTT -LG -DCMut -Dayhoff -WAG -G -I -F -AIC -BIC’) (Darriba et al., 2011; Kalyaanamoorthy et al., 2017; Nguyen et al., 2014). Maximum-likelihood phylogenies were inferred with iqTree using a JTT+I+G4 substitution matrix with USF sequences as outgroups with the following settings: ‘-m JTT+I+G4 −bb 10000 -bnni −nt AUTO -alrt 10000 -abayes’. The resulting tree was finalized using FigTree v1.4.4 (Morariu et al., 2008) and Adobe Illustrator CC 2018.

#### Alignment of Homo sapiens TFEB and Aiptasia MITF-like

Amino acid sequences of *Homo sapiens* TFEB (P19484) and *Aiptasia* MITF-like (XM_020895872.1) were aligned using ClustalW v2.0 (Larkin et al., 2007) in Geneious v10.2.6 (Biomatters).

#### Search for CLEAR elements

*Aiptasia* homologs of human CLEAR network genes (Palmieri et al., 2011) were identified using reciprocal BLAST. Promoters of *Aiptasia* homologs were manually searched for CLEAR elements and E-box elements (palindromic CANNTG) up to 1 kb upstream of the transcription start site in Geneious v10.2.6 (Biomatters).

#### Alignment of SLC38A9 transmembrane domain 8

Amino acid sequences of SLC38A9 homologs in *Homo sapiens* (Q8NBW4), *Danio rerio* (NP_001073468.1), *Xenopus tropicalis* (NP_001011337.1), *Caenorhabditis elegans* (NP_001076680.1), and *Aiptasia* (XP_020892418.1) were analyzed using SMART protein prediction v8.0 (Letunic and Bork, 2017) to identify transmembrane domain 8 (TM8). TM8 sequences were aligned using ClustalW v2.0 (Larkin et al., 2007) in Geneious v10.2.6 (Biomatters).

### Staining procedures in *Aiptasia* larvae

#### Lipid droplet staining with Nile Red

For assessment of maternal lipids in *Aiptasia* larvae, aposymbiotic larvae were fixed 2, 6, and 10 dpf in 4 % formaldehyde for 20 min at room temperature followed by 3 washes in PBS. For assessment of lipid contribution by symbionts, larvae were infected with 10^5^ SSB01 per ml at various time points (2-5 dpf, 6-7 dpf, 8-9 dpf) or left aposymbiotic and fixed 10 dpf in 4 % formaldehyde for 20 min at RT, followed by 3 washes in PBS. Larvae were stained with Nile Red (final concentration 5 μg/ml in 1x PBS from 0.5 mg/ml stock in acetone; N3013, Sigma-Aldrich Co. LLC) for 15 min, followed by staining with Hoechst 33258 (final conc. 10 μg/ml in 1x PBS; B2883, Sigma-Aldrich Co. LLC) for 15 min and 2 washes in 1x PBS before mounting in 87% glycerol in PBS. Images were acquired on a Leica TCS SP8 confocal laser scanning microscope using a 63x glycerol immersion lens (NA 1.30) and Leica LAS X software. Hoechst 33258 and symbiont autofluorescence were excited with the 405 nm laser line, and Nile Red autofluorescence was excited with 488 nm laser. Fluorescence emission was detected at 405 – 480 nm for Hoechst 33258, 540 – 620 nm for Nile Red and 700 – 740 nm for symbiont autofluorescence.

For assessment of lipid contribution by symbionts, the number of lipid droplets in the endodermal tissue was counted. To this end, stacks of whole larvae with a step size of 1 μm were acquired with confocal microscopy. Maximum projections of the center 30 μm were made in FIJI (Schindelin et al., 2012, 2015) using a custom macro (File S2) and the endodermal tissue was manually selected (File S2). In order to measure only signal from host tissue, the signal from symbionts was subtracted from the Nile Red signal. The number of lipid droplets was determined using a custom ImageJ macro (File S3).

#### Lipid droplet staining in polyp macerates

For assessing the abundance of lipids in *Aiptasia* polyps depending on their symbiotic state, aposymbiotic and symbiotic polyps that either had been starved (for 9 days) or fed (3 times per week for 10 days) were macerated and stained with Nile Red. Small polyps (~2 mm oral disc) in 100 μl FASW were pulled through hypodermic needles of decreasing sizes five times each (gauges 23 and 25). The resulting suspension was fixed in 4 % formaldehyde for 20 min and washed in PBS twice before resuspension in 20 μl PBS. 15 μl of tissue suspension was pipetted onto a well of a 10-well PTFE diagnostic slide (631-1371, VWR International GmbH) and left to dry completely. Tissue was rehydrated with 15 μl of MilliQ water before staining with Nile Red (final concentration 5 μg/ml in 1x PBS) for 15 min, followed by staining with Hoechst 33258 (final conc. 10 μg/ml in 1x PBS) for 15 min and 1 wash in PBS before mounting in 4 μl of 87% glycerol in PBS. Images were acquired on a Leica TCS SP8 confocal laser scanning microscope using a 63x glycerol immersion lens (NA 1.30) and Leica LAS X software. Hoechst 33258 and symbiont autofluorescence were excited with the 405 nm laser line, and Nile Red autofluorescence was excited with 488 nm laser. Fluorescence emission was detected at 405 – 480 nm for Hoechst 33258, 540 – 620 nm for Nile Red and 700 – 740 nm for symbiont autofluorescence. Transmitted light from the 488 nm laser was also detected.

#### Cell proliferation assay with EdU

Cell proliferation was determined using the 5-ethynyl-2’-deoxyuridine (EdU)-Click 488 kit (BCK-EDU488, Sigma-Aldrich Co. LLC). Larvae were washed and resuspended in FASW to a density of ~500 larvae per ml. For assessment of cell proliferation in aposymbiotic *Aiptasia* larvae over time, larvae were incubated in 10 μM EdU for 18 h (6 – 24 hpf) or 24 h (24-48 hpf, 48-72 hpf, 120-144 hpf, 144-168 hpf, 168-192 hpf, and 192-216 hpf). For assessment of differences in host cell proliferation between aposymbiotic and symbiotic larvae, larvae were infected at 5 dpf with 10^5^ SSB01 per ml for 24 hours before wash-out. Larvae were incubated with EdU for 24 h periods starting at 5, 6, and 7 dpf and fixed in 3.7 % formaldehyde for 15 min at 6, 7, and 8 dpf, respectively. Following 3 washes in 0.05 % PBS-Tween 20 (PBS-T), larvae were stained with Hoechst 33258 (final conc. 10 μg/ml) for 40 min. Larvae were washed twice in PBS-T and mounted in ~100 % glycerol. Stacks of whole larvae were acquired on a Leica TCS SP8 confocal laser scanning microscope using a 63x glycerol immersion lens (NA 1.30) and Leica LAS X software. Hoechst 33258 and EdU were excited with 405 and 488 nm laser lines, respectively. Fluorescence emission was detected at 410-501 nm for Hoechst 33258 and 501 – 556 nm, for EdU. For enumeration of nuclei and EdU-positive nuclei, pixel classification was performed in ilastik (Sommer et al., 2011) followed by nuclei identification and enumeration in “vision 4d” software (arivis AG) using the blob finder tool.

#### Aiptasia-specific anti-LAMP1 antibody purification

An antibody against the *Aiptasia* LAMP1 homolog (KXJ16564.1) was raised using the peptide IIGRRKSQRGYEKV coupled to the adjuvant keyhole limpet hemocyanin in rabbit (DJ-Diagnostik BioScience, Göttingen). The antibody was affinity purified from the third bleed using the synthetic peptide coupled to N-hydroxysuccinimide esters (NHS)-activated sepharose (17090601, GE Health Care Life Sciences) according to manufacturer’s protocols.

#### Immunofluorescence of LAMP1 and mTOR

Larvae were fixed for 45 minutes in 4% formaldehyde at room temperature (RT), followed by 3 washes in 0.2% Triton X-100 in PBS (PBT) and one wash in PBS. Larvae were then permeabilized in PBT for 1.5 hours at RT, followed by blocking in 5% normal goat serum and 1% BSA in PBT for 1 hour. Primary antibody was diluted 1:100 in blocking buffer (rabbit-α-LAMP1: 1:100; rabbit-α-mTOR (HPA071227, Sigma-Aldrich Co. LLC): 1:12.5, final concentration 4 μg/ml) and incubated overnight at 4°C. After 3 washes in PBT, the secondary antibody (goat-α-rabbit Alexa 488, ab150089, Abcam plc.) diluted 1:500 in blocking buffer was added and incubated for 1.5 hours at RT. Larvae were washed 2 times in PBT, followed by a 15-minute incubation with 10 μg/ml Hoechst 33258 protected from light at RT, and 2 final washes in PBT and 1 in PBS before mounting in 87 % glycerol. Larvae were imaged on a Leica TCS SP8 confocal laser scanning microscope using a 63x glycerol immersion lens (NA 1.30) and Leica LAS X software. Hoechst 33258, Alexa 488, and symbiont autofluorescence were excited with 405, 496, and 633 nm laser lines, respectively. Fluorescence emission was detected at 410-501 nm for Hoechst 33258, 501-541 nm goat-α-rabbit Alexa 488, and 645-741 for symbiont autofluorescence.

### Western blots

#### Western blot analysis of LAMP1 antibody and LAMP1 deglycosylation assay

Two aposymbiotic or symbiotic adult *Aiptasia* were homogenized in 50 mM Tris-HCl pH 7.5, 200 mM NaCl, and 1% NP-40 with 2X Halt Protease Inhibitor Cocktail (78430, Thermo Fisher Scientific) and then sonicated on ice (Sonifier 250, Branson Ultrasonics) with two rounds of 25 pulses at duty cycle 40%, output control 1.8. The homogenate was centrifuged at maximum speed at 4°C for 10 minutes and supernatant was transferred to a new tube. Deglycosylation assay using PNGase F (P0704S, New England BioLabs Inc.) was performed according to manufacturer’s protocol with the exception of incubating the reaction for 3 hours at 37°C followed by overnight at RT. 0.5 mg/ml of LAMP1 antibody was pre-adsorbed with 1 mg/ml of LAMP1 peptide in 4% milk in 0.1% PBT overnight at 4ºC. Untreated and treated extracts were diluted 1:1 in 5X loading dye and heated to 100°C or 60°C, respectively, for 5 minutes. Samples were loaded into a 4-20% precast gel (4561095, Bio-Rad Laboratories Inc.), which was run at 90 V for 15 minutes then at 200 V for 1 hour at room temperature in 1X SDS running buffer. The proteins were transferred onto a nitrocellulose membrane at 0.37 A for 1 hour and 15 minutes at RT in 1X transfer buffer (20% v/v methanol, 200 mM glycine, 25 mM Trizma in water). The membrane was blocked for 1 hour at room temperature in 4% milk in 0.1% Triton X-100 in PBS. The blot was divided in two and incubated in either LAMP1 antibody diluted 1:2000 in blocking buffer or pre-adsorbed LAMP1 overnight at 4°C. The following day the blots were washed in 0.1% Triton X-100 in PBS 3 x 15 minutes at RT. The secondary antibody, goat-α-rabbit-HRP (Jackson ImmunoResearch), was added at a dilution of 1:5000 for 1 hour at RT, protected from light, followed by 3 x 15 minute washes in either 0.1% Triton X-100 in PBS and one final wash in 1X PBS. The blot was developed using 1:1 ECL (GERPN2232, Sigma-Aldrich) and imaging on ECL Imager (ChemoCam, Intas).

#### Western blot analysis of phospho-4E-BP1 (p4-EBP1)

Symbiotic or aposymbiotic *Aiptasia* polyps, that had been either starved for >3 weeks or fed 3 times weekly for the last 3 weeks, were blotted dry on tissue paper and resuspended in 100 μl of 2x loading dye (120 mM Tris-HCl pH 6.8, 20 % glycerol, 2 % SDS, 40 mM dithiothreitol) and incubated at 95 °C for 10 min. Lysed samples were then put on ice, and sonicated (Sonifier 250, Branson Ultrasonics, with two rounds of 30 and 20 pulses, respectively, at duty cycle 40% and output control 1.8), followed by incubation at 95 °C for 5 min. Samples were then pelleted by centrifugation at 2000 rcf for 30 s at room temperature and total protein concentrations of the supernatant were determined via Bradford Assay, masking SDS interference with α-cyclodextrin (Rabilloud, 2018). Samples were stored at room temperature until loading on gels. 30 μg (fed 1 d before sampling) or 45 μg (fed 3 d before sampling) of protein were loaded per sample.

Samples were run on 12 % SDS gels at 90 V (stacking gel) and then 150 V (resolving gel). The proteins were transferred onto a nitrocellulose membrane at 0.35 A for 1 hour at RT in 1x transfer buffer (25 mM Tris, 200 mM glycine, 20% methanol). The membrane was blocked for 1 hour at room temperature in 5% milk in 0.1% Tween 20 in TBS. The blot was cut at the 25 kDa marker band. The top was incubated with α-tubulin antibody (1:3000; T9026, Sigma-Aldrich Co., LLC), the lower half incubated with the p4-EBP1 primary antibody (1:1000; 2855T, Cell Signaling Technology) overnight at 4°C. The following day the blots were washed in 0.1% Tween 20 in TBS 3 x 5 minutes at RT. The secondary antibody, goat-α-rabbit-HRP (115-035-144, Jackson ImmunoResearch) for p4-EBP1 and goat-α-mouse-HRP for α-tubulin (115-035-044, Jackson ImmunoResearch), was added at a dilution of 1:10000 for 1 hour at RT, followed by 3 x 5 minute washes in 0.1% Tween 20 in TBS. The blot was developed using ECL (GERPN2232, Sigma-Aldrich) and imaging on ECL Imager (ChemoCam, Intas).

## Supplemental Information

**Figure S1. Related to Figure 2D.**
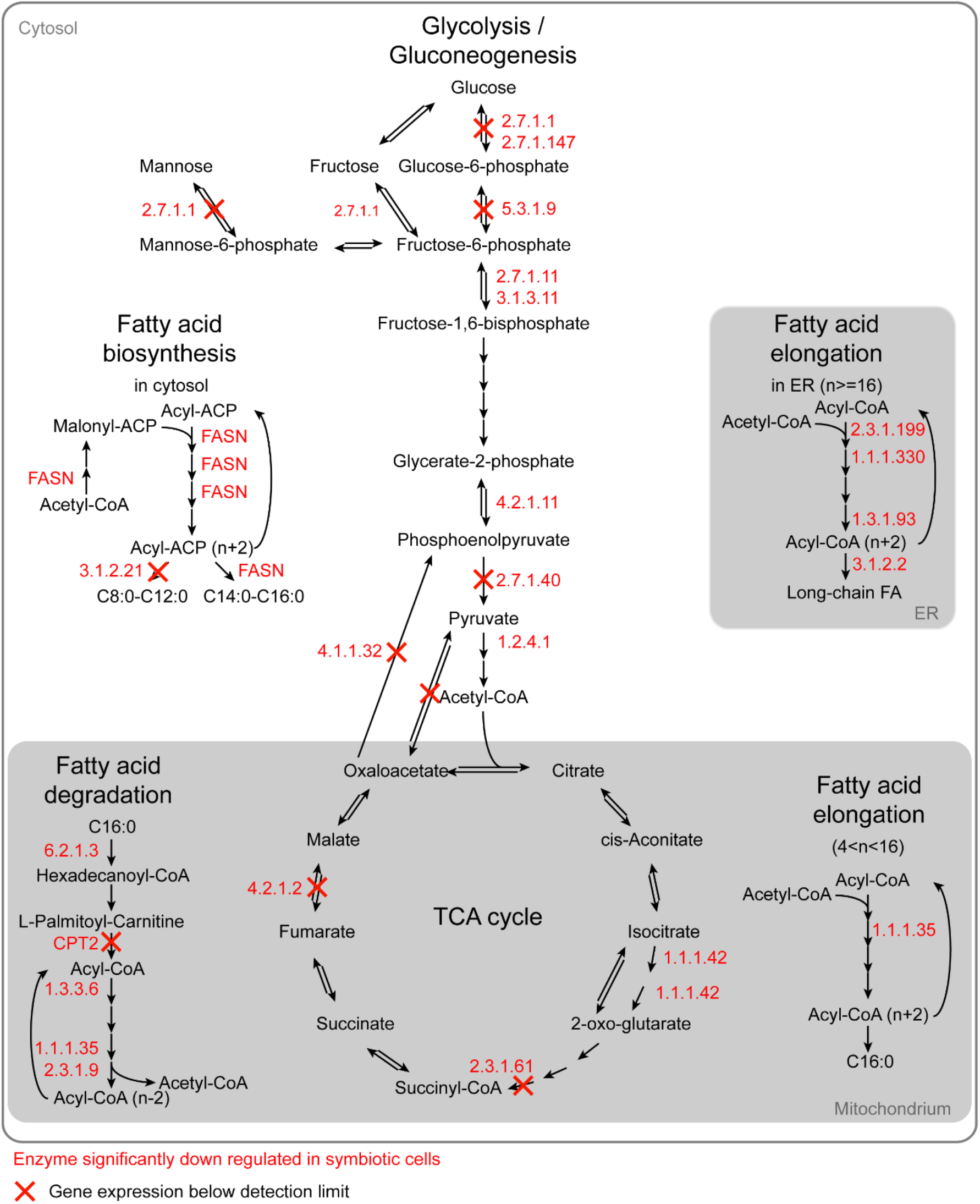
Schematic summarizing gene expression for genes involved in core metabolic pathways based on KEGG pathways. Down-regulated genes are represented by their Enzyme Commission (EC) number in red. When no expression was detected for any gene coding for an enzyme, the corresponding reaction is crossed out in red.

**Figure S2. Related to Figure 3.**
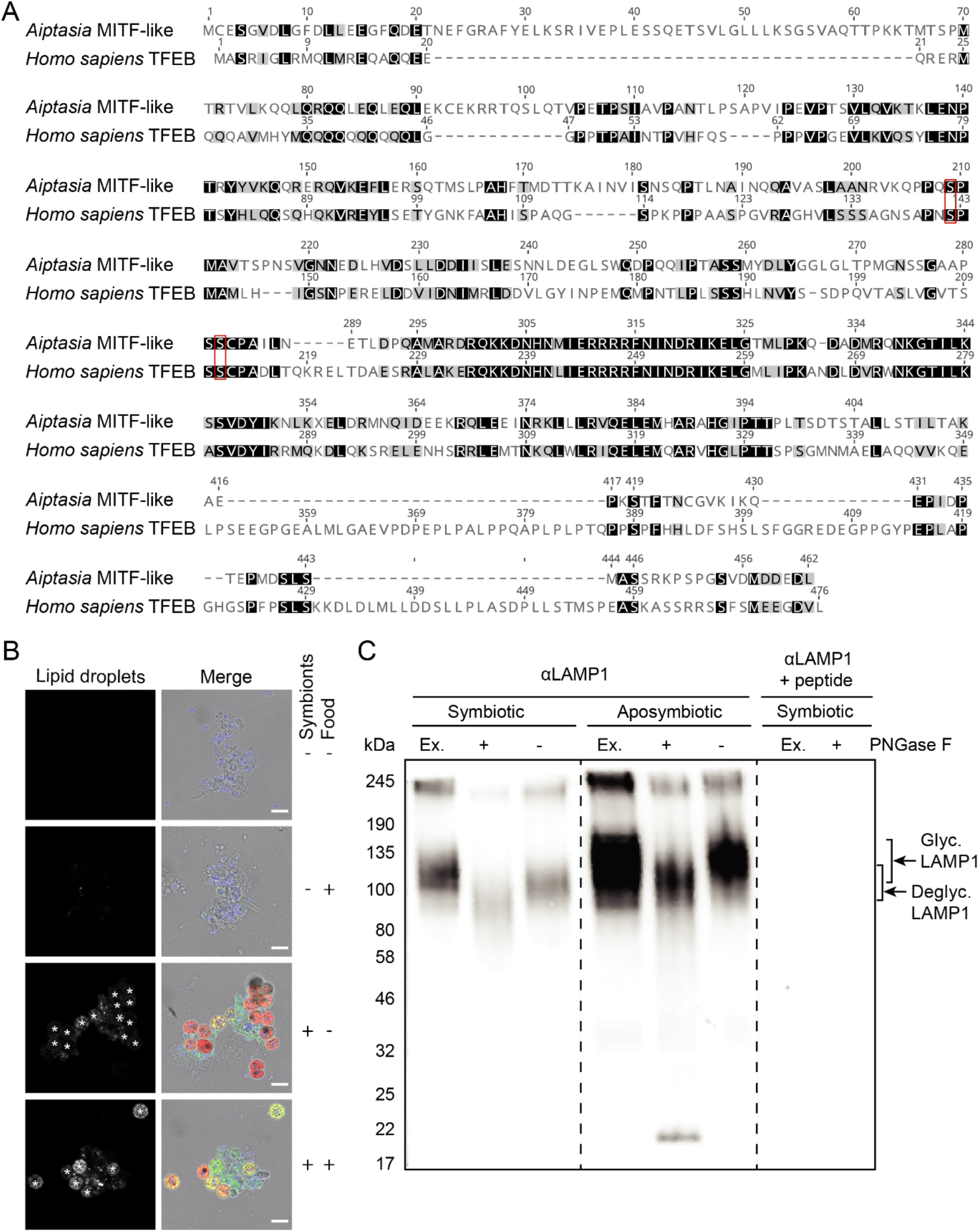
**A)** Related to Figure 3A. Alignment of amino acid sequences of *Homo sapiens* TFEB (P19484) and *Aiptasia* MITF-like (XM_020895872.1). Conserved mTOR-dependent phosphorylation sites are indicated in red. **B)** Related to Figure 3C. Representative images of tissue macerates from *Aiptasia* polyps stained with Nile Red comparing aposymbiotic and symbiotic polyps with food (feeding 3 times per week, for 10 days and fed last two days prior to maceration) and without food (no feeding for 9 days). Asterisks (*) indicate interference of symbiont autofluorescence. Colors in merge are nuclei in blue (Hoechst 33258), lipid droplets in green (Nile Red), symbiont autofluorescence in red, and transmitted light in gray; scale bars represent 10 μm. **C)** Related to Figure 3D. Characterization of α-LAMP1 antibody used in Fig. 3D by Western blot. Due to glycosylation, LAMP1 has been observed to run at a higher than predicted weight (38 kDa) (Winchester, 2001). Accordingly, deglycosylation of homogenates of symbiotic and aposymbiotic adult *Aiptasia* using PNGase F resulted in a shift of the detected LAMP1 signal to a lower molecular weight. In line with its specificity, pre-absorption of the LAMP1 antibody with the peptide used as an antigen abolished signal detection.

**File S1. Related to Figure 1E.**
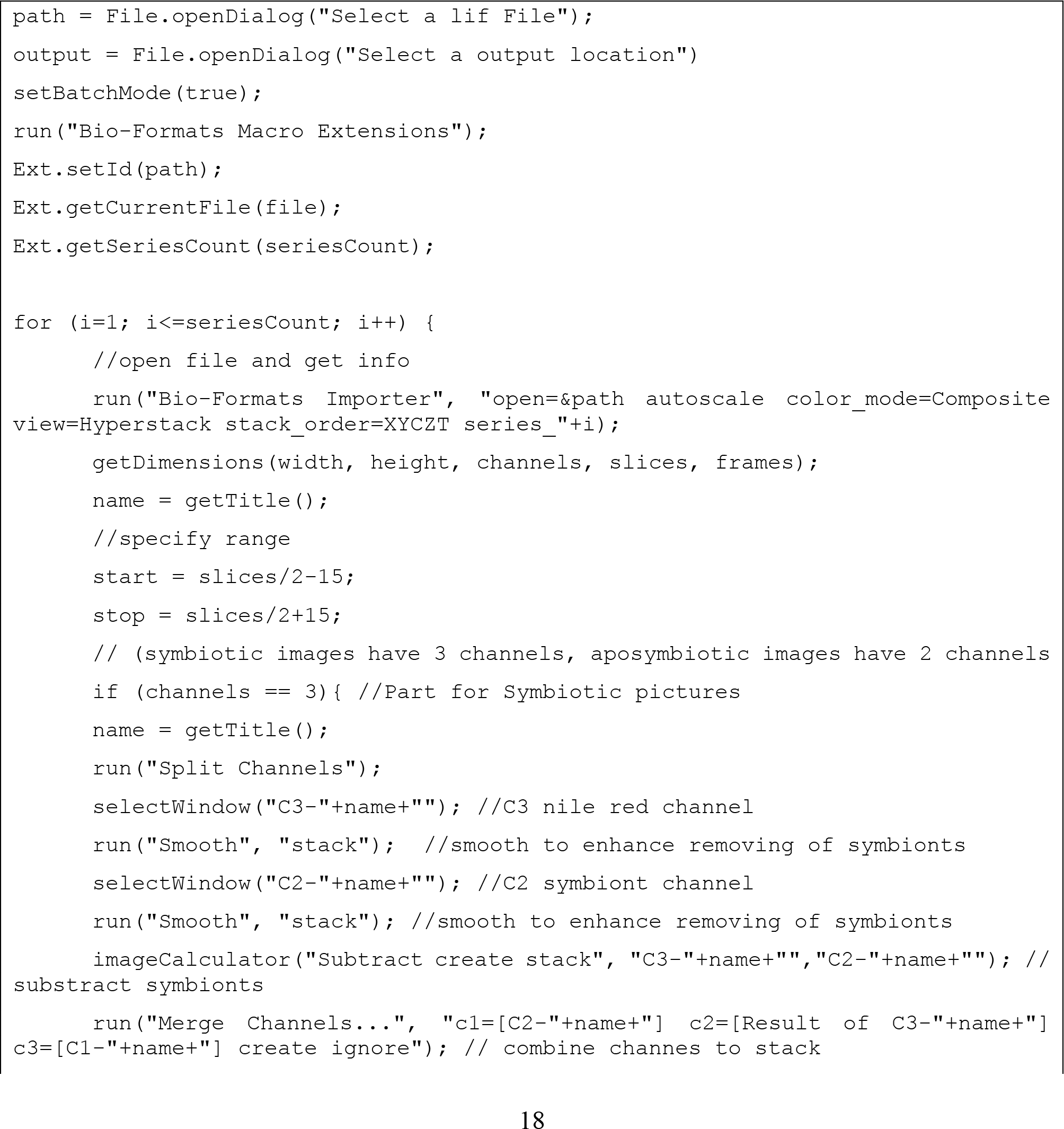
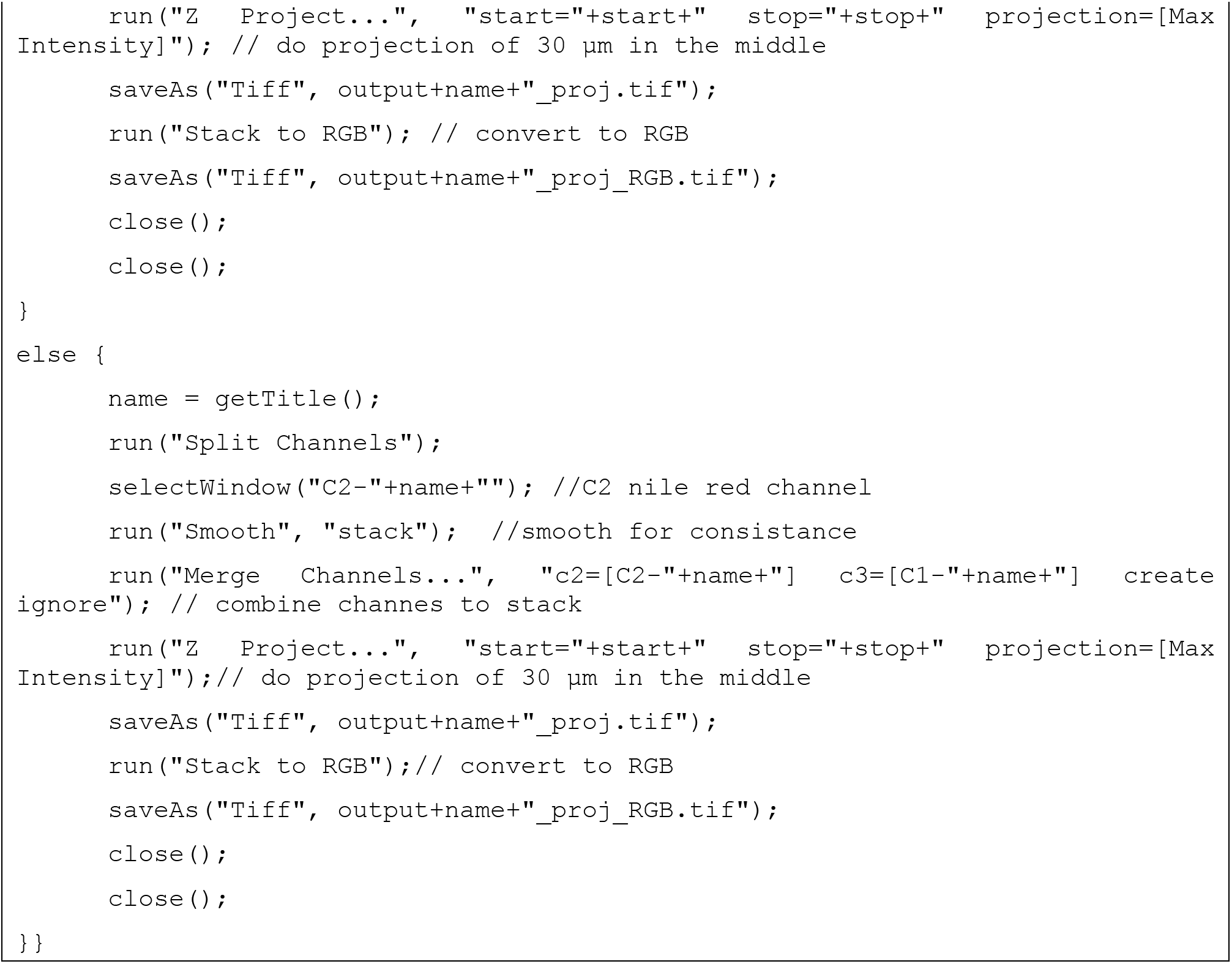
ImageJ macro used to make maximum projections of center 30 μm of Z-stacks.

**File S2. Related to Figure 1E.**
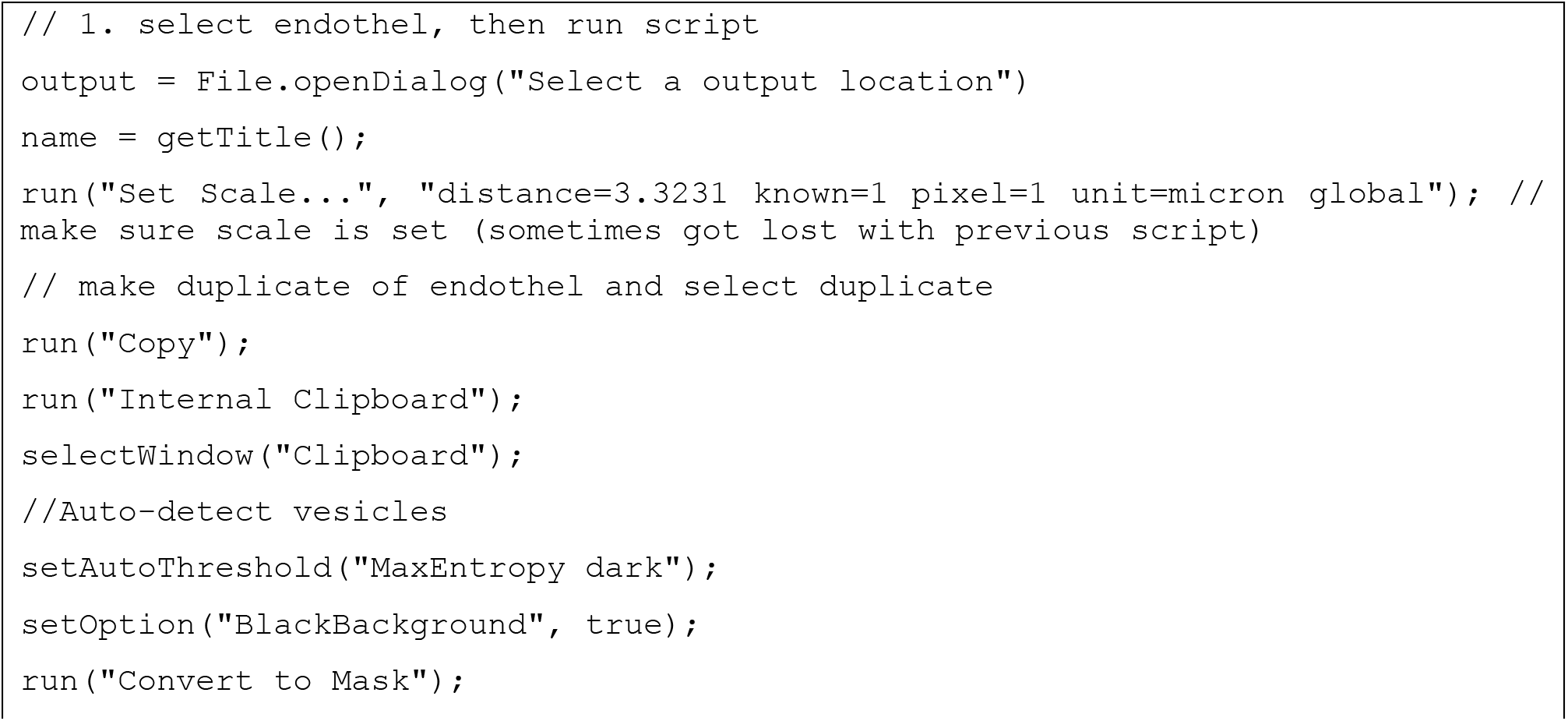
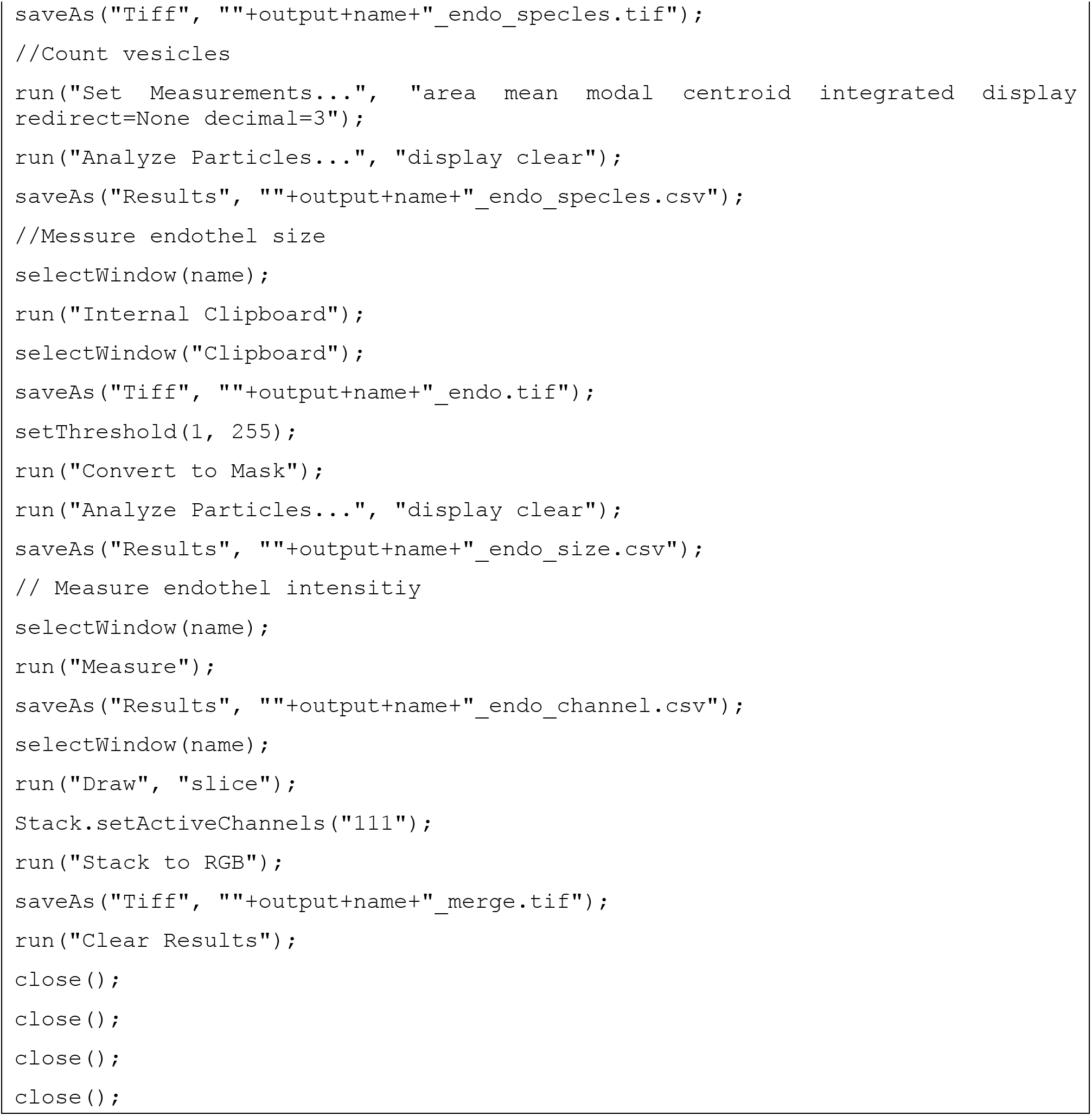
ImageJ macro used to count the number of lipid droplets per in maximum projections.

**Table S1. Related to Figure 2D.** Detailed list containing log2-fold changes between aposymbiotic and symbiotic cells, including TMM values for all replicates. Highlighted cells represent replicates used for differential gene expression analysis with aposymbiotic cells in gray and symbiotic cells in red. Down- and up-regulated genes in symbiotic cells are in different tabs of the .xlsx file.

**Table S2. Related to Table 1.** Enriched KEGG pathways among the down-regulated genes in the symbiotic cells (p-value ≤0.15) including a detailed list of all differentially expressed genes for these pathways.

**File S3. Related to Figure 3A.** Nexus file containing raw sequences of MITF-family genes, trimmed alignments and tree information for the maximum likelihood phylogeny.

